# Engineering genetic circuits in receiver cells for diffusion-based molecular data communications

**DOI:** 10.1101/2023.04.18.536609

**Authors:** Merve Gorkem Durmaz, Neval Tulluk, Recep Deniz Aksoy, H. Birkan Yilmaz, Bill Yang, Anil Wipat, Ali Emre Pusane, Göksel Mısırlı, Tuna Tugcu

**Affiliations:** Department of Computer Engineering, NETLAB, Bogazici University, Istanbul, Turkey; School of Computing, Newcastle University, UK; Department of Electrical and Electronics Engineering, Bogazici University, Istanbul, Turkey; School of Computer Science and Mathematics, Keele University, Staffordshire, UK

**Keywords:** Synthetic biology, molecular communications, genetic circuits, model-driven design, reciever design

## Abstract

Developments in bioengineering and nanotechnology have ignited the research on biological and molecular communication systems. Despite potential benefits, engineering communication systems to carry data signals using biological messenger molecules is challenging. Diffusing molecules may fall behind their schedule to arrive at a receiver, interfering with symbols of subsequent time slots and distorting the signal. Theoretical molecular communication models often focus solely on the characteristics of the communication channel and fail to provide an end-to-end system response, since they assume a simple thresholding process for a receiver cell and overlook how the receiver can detect the incoming distorted molecular signal. There is a need to develop viable end-to-end communication models. In this paper, we present a model-based framework for designing diffusion-based molecular communication systems coupled with synthetic genetic circuits. We describe a novel approach to encode information as a sequence of bits, each transmitted from a sender as a burst of specific number of molecules, control cellular behavior, and minimize cellular signal interference by employing equalization techniques from communication theory. This approach allows the encoding and decoding of data bits efficiently using two different types of molecules that act as the data carrier and the antagonist to cancel out the heavy tail of the former. We also present Period Finder, as a tool to optimize communication parameters, including the number of molecules and symbol duration. This tool facilitates automating the choice of communication parameters and identifying the best communication scenarios that can produce efficient cellular signals.

## 1 Introduction

Cells communicate with the environment and each other to maintain life (*1*). Examples include single-cell organisms, such as bacteria that are organized in micro societies and neurons in the human brain. Naturally, biological and signaling molecules are used as information carriers. The communication mechanisms between sender and receiver cells are of importance for the communications engineers to engineer multicellular applications. Synthetic genetic circuits (*2*, *3*) can offer a rewarding tool for molecular communication systems to go beyond the diffusion of biochemical molecules to modulate and carry information between sender and receiver cells by providing the capability to understand and engineer complex multi-cellular applications. Integrating approaches from molecular communications and synthetic biology and considering intracellular and intercellular dynamics of signaling molecules can help achieve communication-based and robust biological applications. However, the application of such integrative approaches to develop biological communication systems is limited to date.

Molecular communication systems can become challenging to engineer when living cells are involved. Signaling molecules propagating in the intercellular medium may trigger adverse cellular responses (*4*). One way to facilitate the design of predictable biological molecular communication channels is to apply model-based design approaches, which may involve integrating multi-scale mathematical models (*5*, *6*). This process can be complemented by a holistic approach to simulate the dynamics of, and cellular response to, messenger molecules that carry information for signaling and diffuse across a communication channel.

Incorporating the dynamics of underlying transmission channels has several advantages in developing nano- and micro-scale biological communication systems. Inspired by nature, molecular communication systems are bio-compatible and less invasive than their alternatives, such as utilizing electromagnetic radiation (*7*). These systems are energy efficient since the utilized biochemical reactions consume low energy. In addition, the transmitted molecules propagate freely, and molecular communication via diffusion (MCvD) has no external energy requirements (*8*). A well-known model of a molecular communication channel developed by Shannon and Weaver (*9*) consists of five key elements: an information source that generates the message, a sender or transmitter (*Tx*) that encodes the message into a communication signal, a channel in which the signal propagates, a receiver (*Rx*) that decodes or translates the received signal, and a destination node.

Developing signaling channels and encoding information can be affected by the inherent noise and stochastic behavior of diffusing molecules in a crowded medium, such as the intracellular environment full of water, ions, and various macromolecules (*10*). Noise is the undesired element of a communication channel and can be defined as any inference that changes the received signal, typically in a destructive way (*9*). Diffusing molecules may not arrive at the receiver cells on their predestined time slots and may interfere with the subsequent data transmissions (*11*). As a result, when molecules released from a sender represent a particular symbol, the receiver may decode this symbol incorrectly. This situation causes significant intersymbol interference (ISI), which is considered one of the major challenges in diffusion-based communication systems that hinders communication at high data rates (*12*, *13*).

Different theoretical approaches have been proposed to handle ISI (*14*–*16*). Noel and co-workers (*17*) proposed adding enzymes into the propagation channel. Enzymes degrade information molecules and prevent stray molecules from interfering with future transmissions. Tepekule and co-workers (*18*) proposed a molecular transition shift keying technique in which the presence of two different types of molecules (type-A and type-B) is used to encode bit-1. The absence of these molecules represents bit-0. The choice of molecule type to encode bit-1 depends on the following bit-0 or bit-1 symbol. If the next symbol is bit-0, type-B molecules are sent. If the next symbol is bit-1, type-A molecules are sent. This strategy ensures that only type-B molecules are released before bit-0, and the accumulation of molecules and ISI are restrained (*18*). Another proposed solution is the pre-equalization method (*19*), which involves transmitting two different molecule types from a sender: type-A information encoding molecules and type-B destructive molecules. In this approach, type-B molecules eliminate the effect of the stray type-A molecules. The impact of the destructive molecule is imitated by employing a subtraction operation at the receiver. However, the aforementioned methods do not address the issue of how these molecules are converted to intracellular signals that affect the system’s response and noise when genetic receivers are employed. This study incorporates and improves the pre-equalization method that has the reduced complexity compared to other methods to address the effects of intercellular and intracellular signaling processes.

Different modulation techniques have been proposed to encode information using molecular communication systems. Some of these techniques are based on the concentration of the transmitted molecules from a *Tx* sender (*20*), the type of the transmitted molecules (*21*), the release time of the transmitted molecules within a communication time slot (*22*), and the locations of spatially-separated multiple transmitters (*23*). There are also hybrid techniques that combine more than one aspect of the information carriers (*24*, *25*).

The movement of molecules in a molecular communication channel can be modeled by the diffusion process or the Brownian motion. In a fluidic environment without any flow, molecules move randomly (*26*). When diffusing molecules reach receiver cells, they may activate some processes or yield information bits after a demodulation process. Therefore, evaluating the expected number of received molecules is critical for designing an effective MCvD system that involves receiver cells. Channel properties, absorption rate, and biochemical kinetic rates may affect the activation of a synthetic genetic circuit inside an *Rx* receiver. The activation of the circuit may also depend on the specific level of intercellular signaling molecules for a certain amount of time. It is desirable to incorporate models of genetic circuits to design efficient molecular communication systems and understand the effect of ISI on cellular response.

Various modeling formalisms exist to analyze the dynamic behavior of genetic circuits. For example, the Systems Biology Markup Language (SBML) (*27*) standardizes the representation of biochemical reactions. Tools such as COPASI (*28*) can simulate SBML models deterministically or stochastically to gain insight into emerging cellular behavior. Such modeling formalisms have already been used to demonstrate the model-driven design of synthetic genetic circuits. The Virtual Parts Repository (VPR) (*29*, *30*) provides reusable, modular, and mathematical models of biological components such as promoters, ribosomal binding sites (RBSs), and coding sequences (CDSs). These models can be joined together to create models of desired systems. The resulting models are hierarchical, based on models of parts and interactions and their aggregation. This model-driven approach is ideal for designing and optimizing genetic circuits and computationally exploring large design spaces of desirable biological systems.

Standardization efforts are essential to exchange information between tools without loss of data. Synthetic genetic regulatory circuits can be computationally represented using the Synthetic Biology Open Language (SBOL) (*31*, *32*). SBOL designs can include constraints to capture the sensing of external molecules and descriptions of intended biochemical reactions. This qualitative information can consequently be used to create models that can be simulated. The recent version of VPR provides SBOL-to-SBML conversion to automate the generation of computational models of genetic circuits (*30*, *33*).

Here, we present a computational modeling approach to facilitate the design of molecular communication systems that can be coupled with living cells. This approach is workflow based and integrates the modeling efforts in molecular communications and synthetic biology. Computational simulations are used to design different communication aspects to encode and send information using biological molecules. Our modeling framework aims to minimize signal interference and allows computationally optimizing key communication parameters, such as symbol duration, to produce efficient cellular signals.

## 2 Results

The computational modeling approach presented here was developed to design communication systems using molecular and biological communication channels. This process involves coupling intracellular and extracellular processes with diffusion dynamics and three-dimensional molecular channel propagation (*34*). The information can be encoded as sequential bits, each representing a number of molecules released by a sender. As a result, a response signal is created at a receiver via the accumulation of cellular molecules. We demonstrate this approach computationally using a receiver design based on synthetic and bacterial genetic regulatory networks to decode information. The coupling of intercellular and intracellular mechanisms is implemented as a workflow in which an MCvD system with pre-equalizer (*19*) can eliminate ISI. The workflow incorporates VPR to facilitate model-driven design and optimization of genetic receivers.

### A Pre-equalizer for Engineered Receiver Cells

ISI mitigation is especially challenging when living cells act as receivers due to different timescales in intercellular diffusion dynamics and intracellular biochemical reactions. Different biochemical rate parameters may cause received signals to interfere with the subsequent cellular signals. In this work, we propose a cellular pre-equalizer method building upon the literature (*11*). This method involves two input signals emitted at a sender and two additional cellular signals at a receiver.

The input signals together carry a single bit of data to reduce interference. The first signal (type-A) is the actual data carrier, and each bit corresponds to type-A molecules being sent over a specific period. The second input signal (type-B) removes the heavy tail of the type-A signal. The effectiveness of the removal of the interference is calculated by the receiver cells. Hence, *A_out_* and *B_out_* signals are transformed into intracellular molecules *A_in_* and *B_in_*, respectively. *A_in_* is the observable molecule that relays the signal, and *B_in_* is the antagonist of *A_in_* to cancel out the right amount of *A_in_* to mitigate the adverse effects of ISI.

It is crucial to comply with the processing rates of *Rx* receiver cells to send sequential information or data bits. This process requires maintaining a specific level of messenger molecule concentration. Moreover, to eliminate the heavy tail of *A_in_* at an *Rx* receiver, *B_out_* is emitted *t*_shift_ seconds after *A_out_* is released at a *Tx* sender. With the appropriate communication parameters, the deteriorating effects of the ISI molecules can be eliminated by considering the difference between the two cellular molecule types (*A_in_ − B_in_*). Hence, communication performance can be improved significantly. However, determining the parameters of the pre-equalizer, exploring potential communication scenarios, and measuring the performance of each scenario’s end-to-end communication performance is not trivial. This design space exploration process is carried out by Period Finder.

### Communication Period Finder

A tool called Period Finder has been developed to implement the proposed pre-equalizer method and efficiently design communication channels. Period Finder evaluates the communication performance and optimizes pre-equalizer’s parameters to minimize the degrading effects of ISI. These processes are achieved using computational simulations.

Period Finder generates possible signal propagation scenarios with different signal parameters with an effective *A_out_*/*B_out_* ratio to mitigate ISI. The resulting scenarios are evaluated to ensure that each bit persists for the desired period or symbol duration. The signal parameters of these scenarios generated by Period Finder (Figure 2) are assessed through performance metrics involving MOL-eye diagrams (*35*). MOL-eye is similar to the ‘eye’ diagram that is used for measuring the quality of signals in conventional communication schemes and is adapted to molecular communications. As a result, each communication scenario is scored, and the most effective scenario with the highest score in terms of *A_out_*/*B_out_* is selected. This approach allows adjusting the amplitude and symbol duration ratio of the actual data carrier signal to that of the antagonist signal. Consequently, ISI is minimized.

**Figure 1:**
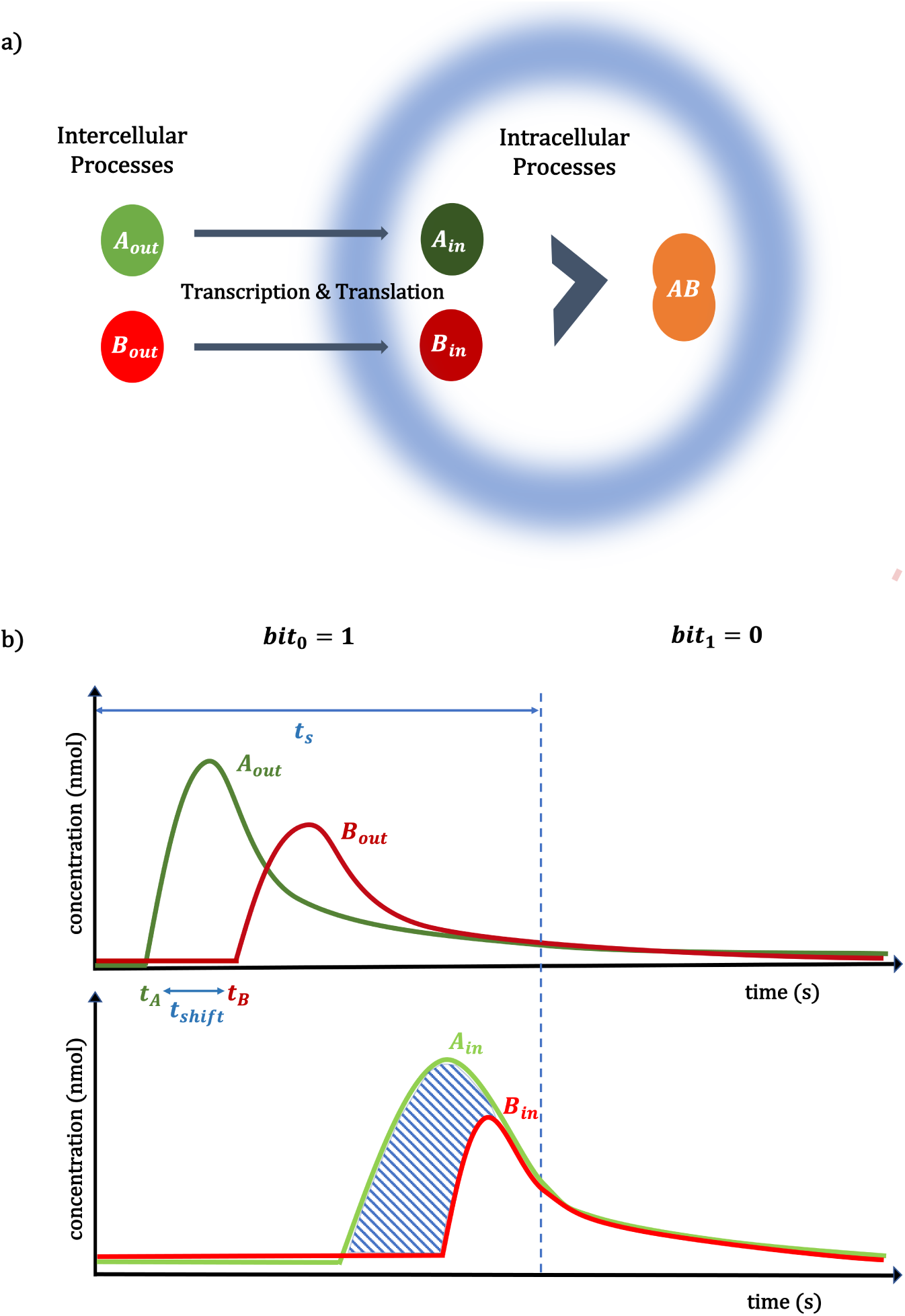
(a) The schematic representation of the general framework for Period Finder, where intercellular signals are converted to intracellular signals. (b) A hypothetical illustration of the signals, where bit-0 represents no transmission and bit-1 represents the transmission of the molecules. In the upper graph, *t_A_* represents the transmission time of *A_out_*, and *t_B_* represents the transmission time of *B_out_*. *t_B_*-*t_A_* is *t_shift_*, and *t_s_* denotes the symbol duration. *A_out_*molecules are transmitted into intercellular space. This signal is converted into intracellular *A_in_* signal with a delay due to cellular processes such as transcription and translation. *B_in_*molecules mitigate the heavy tail of the *A_in_* signal. *A_in_*-*B_in_* is denoted by the shaded region.

**Figure 2:**
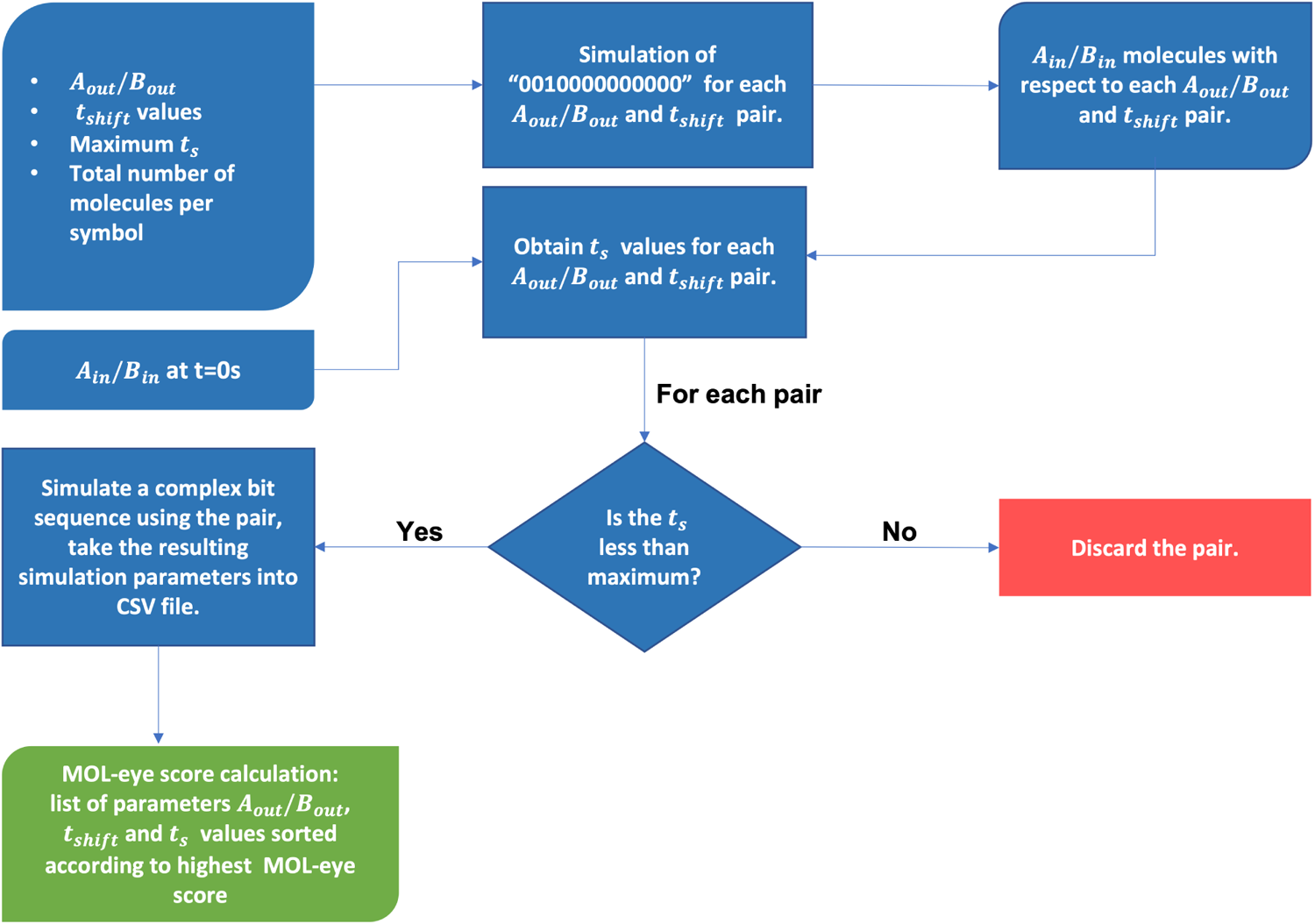
Flowchart of Period Finder. The input of the program is a CSV file containing different *A_out_*/*B_out_* ratios at different *t_shift_* values, a desired maximum *t_s_* value, total number of molecules per symbol, and *A_in_*/*B_in_* before any transmission (t=0). For each *A_out_*/*B_out_* and *t_shift_* pair, the “0010000000000” bit sequence is simulated, and a ratio of *A_in_*/*B_in_* is obtained. Values of *t_s_* are obtained for each resulting *A_in_*/*B_in_*. If the resulting *t_s_* values are larger than the maximum *t_s_* value, they are discarded. Otherwise, a complex bit sequence is simulated using the pair, and the resulting parameters are written into a new CSV file to be evaluated by MOL-eye calculation. After the analysis, *A_out_*/*B_out_* and *t_shift_* pair and *t_s_* values are sorted according to the highest MOL-eye score.

The flowchart of the program is shown in Figure 2. Period Finder retrieves the distinct periods (*t_s_* values) that take the system to return to equilibrium, which is the base state of the cell before any symbol is received. A vector of different *A_out_*/*B_out_* values coupled with a vector of *t_shift_* values, the desired maximum *t_s_* to limit the symbol duration retrieved, and the total number of molecules sent per symbol are given as input. The initial state of *A_in_*/*B_in_* in a receiver cell (*R_x_*) without any inputs is shown in Figure 3a. After the warm-up period (first two bits), the system reaches an equilibrium state where *A_in_*and *B_in_*decrease until they converge on a steady state. Period Finder then creates diverse scenarios with the vectors of *A_out_*/*B_out_* values coupled with a vector of *t_shift_* values to simulate “00100000000000”, a one-shot signal where bit-0 represents nothing is sent, and bit-1 represents that *A_out_* and

**Figure 3:**
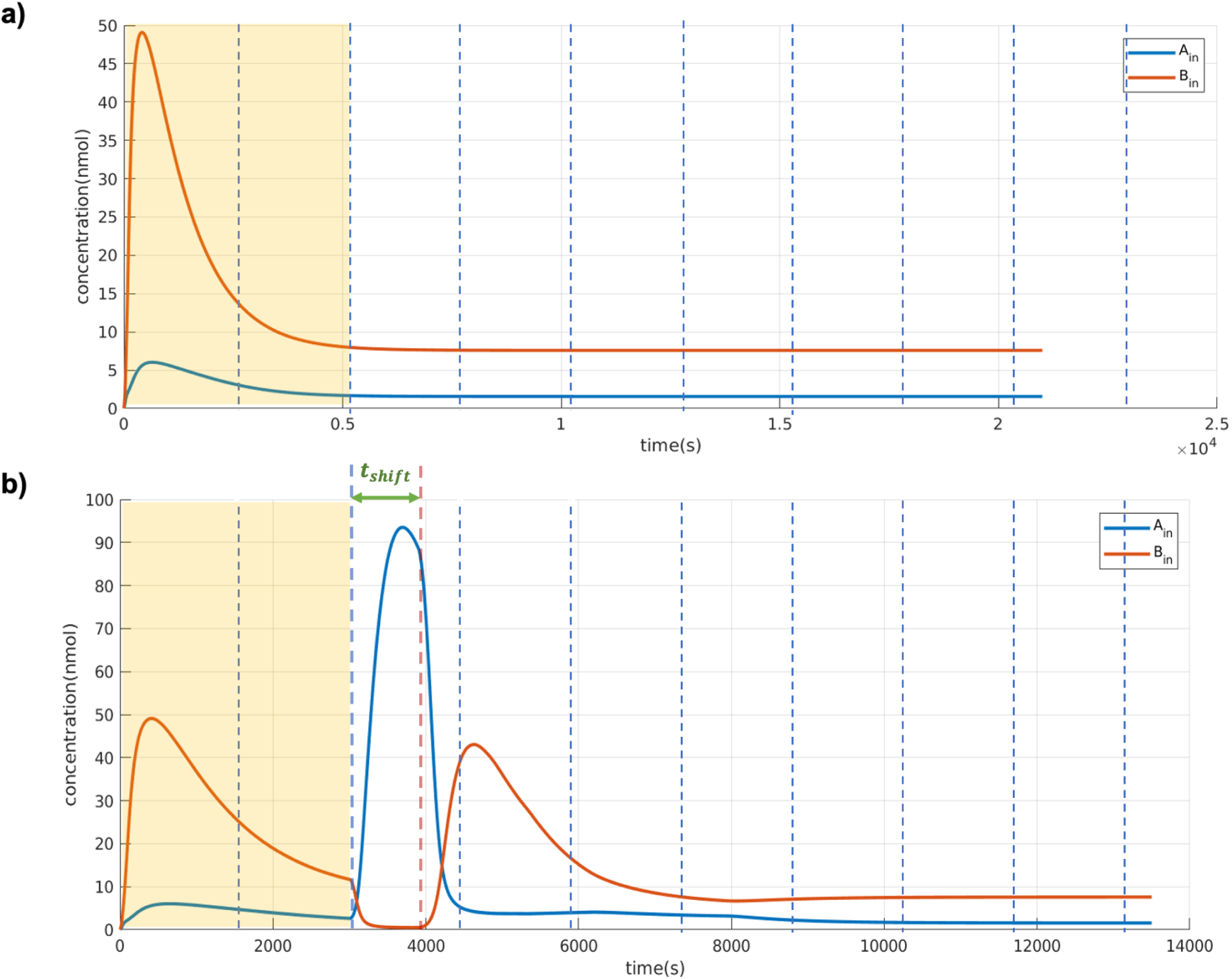
Required inputs for Period Finder (a) Native state of *Rx*, (b) *Rx* response for a one-shot signal of both *A_out_* and *B_out_*. There is a *t_shift_* time between the transmission of *A_out_* and *B_out_* to keep the *A_in_* signal as large as possible in a pre-defined *t_s_*. Simulation parameters are given in Table 1. The first two *t_s_* symbol durations are marked with yellow to indicate the warm-up period.

**Table 1:**
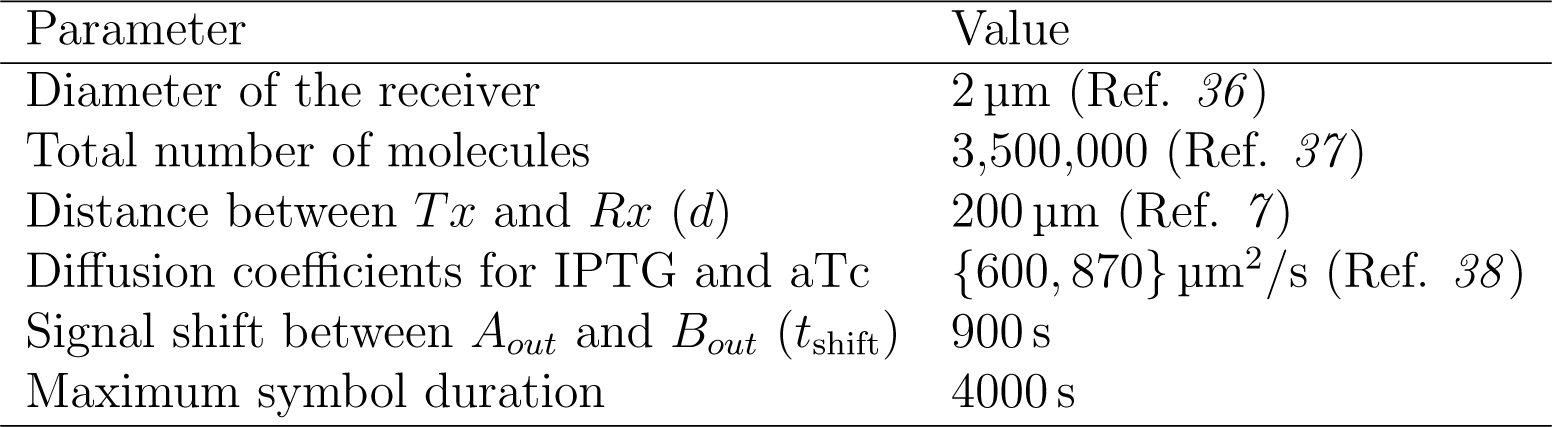
Simulation parameters and values

| Parameter | Value |
| --- | --- |
| Diameter of the receiver | 2 $\mu\text{m}$ (Ref. 36) |
| Total number of molecules | 3,500,000 (Ref. 37) |
| Distance between $Tx$ and $Rx$ ($d$) | 200 $\mu\text{m}$ (Ref. 7) |
| Diffusion coefficients for IPTG and aTc | {600, 870} $\mu\text{m}^2/\text{s}$ (Ref. 38) |
| Signal shift between $A_{out}$ and $B_{out}$ ($t_{shift}$) | 900 s |
| Maximum symbol duration | 4000 s |

*B_out_* molecules are sent. In the one-shot signal, bit-1 is sent only once in the sequence shown in Figure 3b. After *A_in_* and *B_in_* reach their maximum levels in the simulation, the program obtains the corresponding symbol duration to return to equilibrium. Here, we send enough bits to observe the system’s behavior and simulate the scenario in a limited time. If the achieved *t_s_* value is larger than the desired maximum *t_s_* value, then it is discarded. Waiting for the equilibrium state to be achieved ensures the system is on hold until the channel is cleared out to send the subsequent symbol, decreasing the effect of ISI.

Simulation results are stored in a CSV table that includes the *A_in_*/*B_in_* values of oneshot signals for every *t_shift_* and *A_out_*/*B_out_* ratio pair and their corresponding *t_s_* values. The simulation results are evaluated according to MOL-eye diagram performance metrics, and the scenarios with the highest MOL-eye score is reported.

### Minimizing Interference in MCvD

A biological subtraction operation can minimize cellular interface when two signals are used. Here, this operation has been implemented using a design pattern involving two molecules that can bind together (*39*). Inside a receiver cell, *A_out_* activates the production of *A_in_*, and *B_out_* activates the production of *B_in_*. The difference between *A_in_* and *B_in_* molecules is evaluated via a biological subtraction operation. In this design, *B_in_* sequesters *A_in_*. The remaining *A_in_* molecules represent the result of the subtraction operation. Conversion of an intercellular signal to an intracellular signal can be achieved in multiple ways. This process involves designing and incorporating genetic circuits. Here, we adopt transcriptional activation and inhibition processes to control the production of proteins corresponding to different signals. Interactions of these proteins form the basis of our biological program.

In this design, IPTG and aTc diffusing molecules represent the *A_out_* and *B_out_* signals (Figure 4). LacI and TetR repressors inhibit the production of *A_in_*and *B_in_*, respectively. Hence, IPTG activates the production of *A_in_* by inhibiting LacI, and aTc activates the production of *B_in_* by inhibiting TetR. Molecules, such as ExsD and ExsA, that can bind together represent the *A_in_* and *B_in_* molecules (*40*, *41*). Here, ExsD and ExsA were chosen since they can act as TFs to control subsequent cellular response.

**Figure 4:**
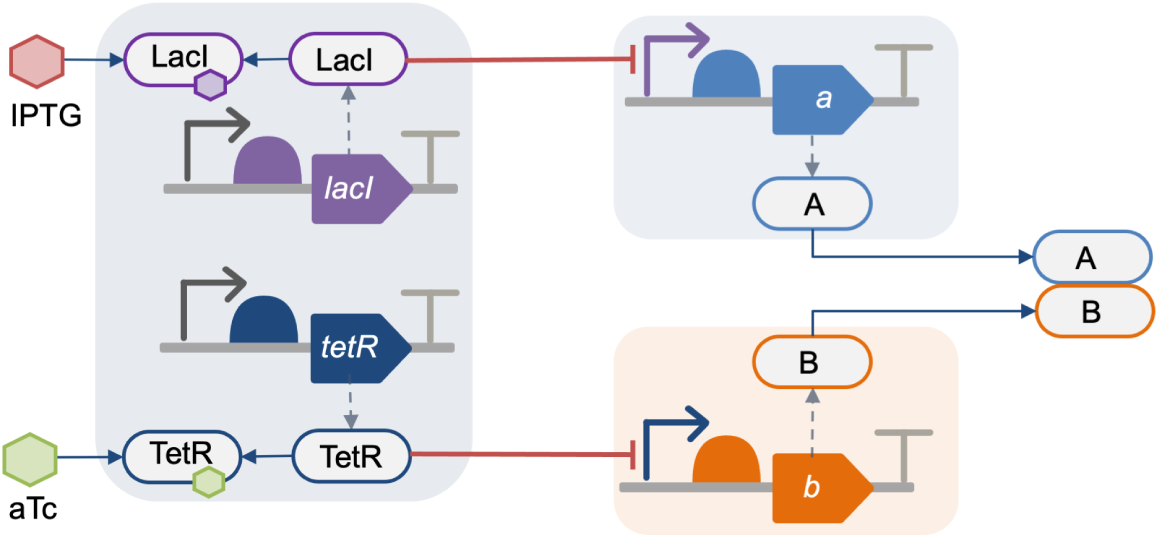
Representation of the system model used in simulations to perform the biological subtraction operation. Remaining *A_in_* molecules over a defined period represent a data bit.

Period Finder relies on VPR to create models of genetic circuits that receive the external *A_out_* and *B_out_* signals and convert them into cellular *A_in_* and *B_in_* signals (Figure 1a). Models of genetic parts such as promoters, RBSs, CDSs, and their interactions are joined together to derive SBML models that can be simulated. Each resulting SBML model represents a possible solution and is evaluated by Period Finder. Models include rate parameters for various biological processes involving genetic production, complex formation, and binding. Period Finder uses the COPASI engine to automate the simulation of each model and determines *A_in_* and *B_in_* values according to the simulation results.

Simulation results when the *B_out_* pre-equalizer is not incorporated are shown in Figure 5. The bit sequence for this simulation is 0010111100101, and the simulation parameters from Table 1 are applied. The first two symbol duration slots represent the warm-up period (*42*) for which the system is unstable. Hence, the simulation starts after this warm-up period. The first two bits are always chosen to be 00 to provide enough time for the system’s stabilization. The lack of the pre-equalizer causes ISI and the accumulation of stray molecules. As shown on the graph, after the 5th symbol, consecutive bit-1 symbols cause the accumulation of molecules. Consequently, the concentration of signal molecules at the 9th symbol of bit-0 is misleadingly higher than expected.

**Figure 5:**
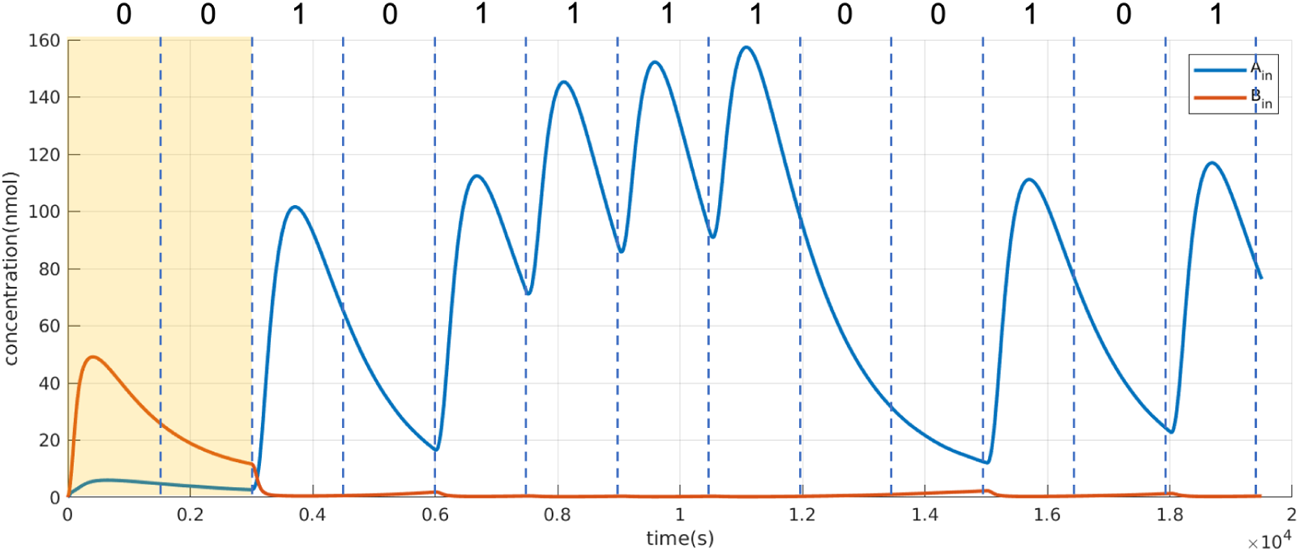
*Rx* response without using the *B_out_* pre-equalizer. The bit sequence of the signal (0010111100101) is shown in the upper part of the graph. The first two *t_s_* symbol duration slots are marked with yellow color to indicate the warm-up period, the y-axis shows the concentration (nmol) of molecules *A_in_* (blue line) and *B_in_* (red line), and the x-axis shows the time in seconds. After a train of consecutive bit-1 symbols, the signaling molecules accumulate in the channel. The concentration of signaling molecules at the 9th symbol of bit-0 is misleadingly high due to the effect of ISI, which is likely to cause an incorrect detection at the receiver side.

Period Finder applies the pre-equalizer approach and adjusts the ratio of the amplitudes of *A_out_* and *B_out_* signals and their symbol duration to decrease ISI. As a result, *B_in_* molecules bind and reduce the number of stray *A_in_* molecules remaining after the symbol duration.

Period Finder evaluates the resulting scenarios according to MOL-eye performance metrics and reports the optimum communication scenarios with the highest MOL-eye scores. An example, MOL-eye diagram of the optimum scenario is shown in Figure 6 where the subsequent signals are superimposed to a single composite graph and the area between the minimum of bit-1 symbol and maximum of bit-0 symbol is denoted as eye-height. The noise can be expected to be the highest when eye is in the most closed form, where it becomes challenging to distinguish bit-1 and bit-0 symbols. However, considering the optimum communication scenarios identified by Period Finder, even when the minimum of bit-1 and maximum of bit-0 is used, the eye-opening is still clear as shown in Figure 6b.

**Figure 6:**
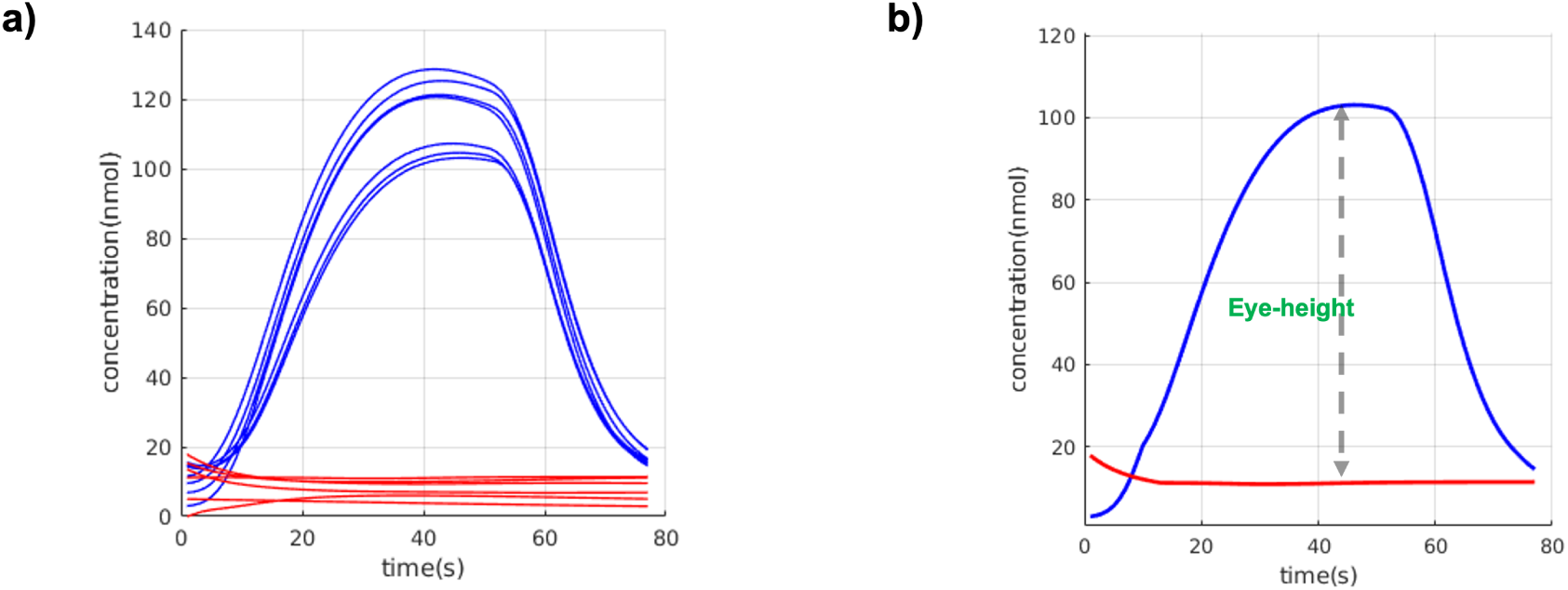
MOL-eye diagram, where successive bit-1 and bit-0s are superimposed, red lines represent bit-0 and blue lines represent bit-1. (a) Overlapped representation of bit-1 and bit-0 symbols (b) The eye-height is the vertical opening between the maximum of bit-0 and minimum of bit-1.

The same communication scenario is shown in Figure 7, including simulation results for both *A_in_*and *B_in_*signals in more detail. As expected, when the 0010111100101 bit sequence is sent, consecutive bit-1 symbols in the middle do not cause the accumulation of the signaling molecules anymore. Moreover, each bit-1 and bit-0 information is exchanged clearly, demonstrating that the effects of ISI are drastically reduced.

**Figure 7:**
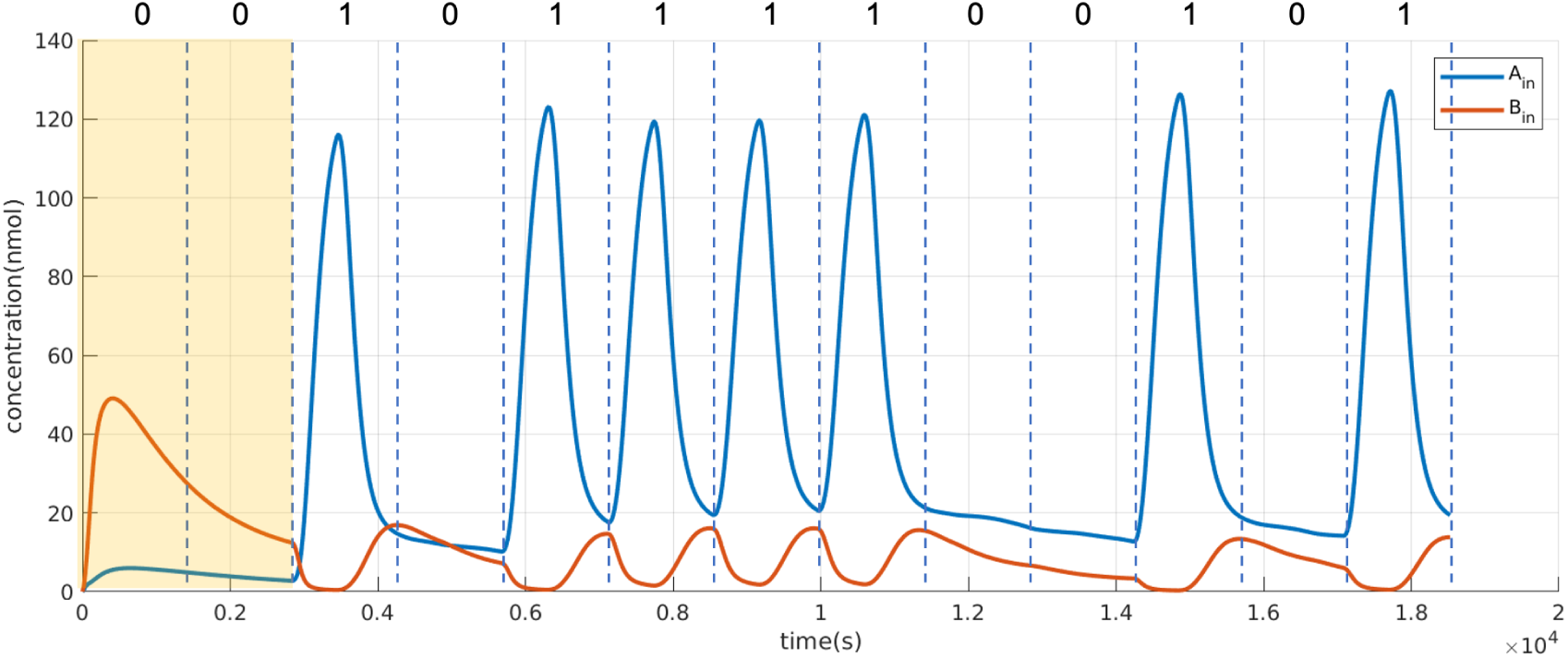
*R*x response using the *B_out_* pre-equalizer. *A_out_*/*B_out_* is 0.15. The first two *t_s_* symbol durations are marked with yellow to indicate the warm-up period. The solid blue and red lines show the concentration of *A_in_* and *B_in_* molecules over time, respectively. As desired, signaling molecules do not start accumulating during the 5th symbol duration. Furthermore, the concentration of signaling molecules at the 9th symbol of bit-0 is as expected since the pre-equalizer (*B_in_*) mitigates the effect of ISI.

## Discussion

Molecular communication systems have several advantages in developing biological applications. However, it is challenging to develop such applications, which involve encoding and decoding information due to the noisy and uncertain environment of a medium. When these communication systems are coupled with cells that act as receivers, this complexity increases even further due to the large number of genetic parts that can be chosen to decode information and control cellular response. Trial and error-based wetlab experiments can be costly. Moreover, wet-lab experimentation is out of reach for most researchers in the communication engineering area, since it requires qualified laboratories, equipment, and specialized staff (*43*). Simulation environments can provide valuable insights to overcome these issues and develop and test novel communication models. However, this process is not trivial. Diffusion-based cellular processes can be complex and may have different timescales. Resulting computational models may provide insights with varying details for different processes. The work presented here involves algorithms, design patterns, and a simulation approach to overcome the obstacles in engineering receiver cells that function via molecular communications and diffusion of molecules to encode and send information.

One of the main challenges in using engineered receiver cells and diffusion-based systems is decoding the information due to inherent noise. Our work extends the previously proposed pre-equalizer approach (*18*), which involves using two different signals to distinguish data bits, by incorporating two additional cellular signals. A biological subtraction operation for these cellular signals has been defined as a genetic circuit design to improve the molecular channel response, reduce cellular noise and control cellular response.

Period Finder can propose successful communication scenarios based on the transmission time difference between the signaling molecules and their concentration ratio. These scenarios are ranked using the MOL-eye performance metrics. Hence, Period Finder can be ideal to automate exploring different communication parameters via computational simulations.

Nature-inspired communication systems can have several advantages for developing biological applications. On the other hand, it may be challenging to implement expectations of generic communication systems. Notably, a communication scenario can be slow. Here, we explored minimizing symbol interface by considering symbol duration, which can be minutes due to the diffusion of molecules and accumulation of cellular molecules via transcription and translation events (*44*). As a result, a communication scenario involving a series of data bits may take hours.

Another challenge in our experiments is establishing the warm-up period. In a discrete-event simulations, the system is initially empty and idle, and the steady state has not been reached. The situation is different in biological systems, due to the leakiness and basal expression of biological molecules, which can cause inaccurate results (*45*). To improve a molecular channel’s efficiency, decoding information in receiver cells should start after a sufficient warm-up period for the system to reach the initial steady state. In our simulations, this steady state is reached after the first two symbol durations. Hence, the first two symbols are chosen as bit-0 in our simulations. Results within the warm-up period are ignored to improve the accuracy of overall results.

The modeling approach presented here demonstrates how efforts in molecular communications and synthetic biology can be combined to provide an integrated view of intracellular and intercellular processes to design novel communication systems. Our approach allows validating molecular communication designs *in silico* and identifing suitable system parameters computationally to inform wet-lab experiments.

## 3 Methodology

Period Finder was implemented in MATLAB and Java to produce efficient data signals and mitigate ISI caused by diffusion and cellular noise. Diffusion models were integrated with cellular models in a multi-scale approach, and simulations were automated. Our models incorporate cellular signaling to integrate the effect of diffusing molecules to trigger a desired cellular activity within an *Rx* receiver, for example, to convert a molecular signal into a cellular signal. Here, we designed a genetic circuit that builds upon gene regulatory networks involving transcriptional activation and repression processes to control cellular output in response to sensing intercellular (*A_out_* and *B_out_*) signaling molecules. Molecular interactions between different circuit components are used to evaluate intracellular signals (*A_in_ −B_in_*).

Diffusion dynamics were linked to cellular genetic circuit models via molecular communication parameters. These parameters were used to derive genetic circuit models, each representing a possible design. Genetic design automation was implemented in Java. VPR2 (*30*) was used to define models of biological parts such as promoters, RBSs, and CDSs and molecular interactions, such as TF-promoter activation and inhibition, complex formation, and degradation. The model construction process was facilitated using the SVPWrite language to specify the order and types of biological parts. For example, the “prom1:prom;rbs1: rbs;cds1:cds;ter1:ter” input specifies a single transcriptional unit where “prom1” is a promoter, “rbs1” is an RBS; “cds1” is a CDS and “ter1” is a terminator. The resulting genetic circuit definition was represented in SBOL2(*31*), and information about molecular interactions was added. Hierarchical SBML models (*46*) representing potential genetic circuits were derived via VPR2’s SBOL-to-SBML conversion. The SBML models were simulated using the COPASI Java bindings (*28*). The simulation results were evaluated by Period Finder. The MOL-eye diagram is adopted for the evaluation of potential communication scenarios.

The movement of particles in a three-dimensional space can be represented via three independent displacements, one for each dimension, where each displacement follows a normal distribution with zero mean and *σ*^2^ variance, denoted as

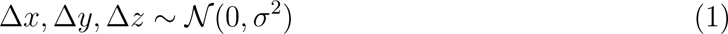

where *t* is the time, 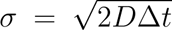, and *D* is the diffusion coefficient that describes the molecule’s mobility inside the fluid (*26*).

Assuming a simple MCvD channel without flow, the expected fraction of diffusing type-A molecules which will reach and be absorbed by the *Rx* during the time frame *t_k_* can be calculated as

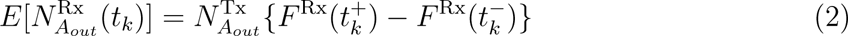

where *E*[·] is the expectation operator, 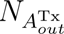 is the number of emitted molecules, *F*^Rx^(*t*) is the time-dependent formula for the expected cumulative fraction of arriving molecules (*34*), 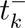 is the start and 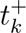 is the end of time frame *t_k_* (*18*). For a simple and symmetric topology like a point transmitter and a single spherical absorber, *F* ^Rx^(*t*) is known analytically(*34*).

In the general model shown in Figure 1, the resulting proteins inside the *Rx* (*A_in_* and *B_in_*, respectively) can bind together. Therefore, *B_in_* can eliminate the effect of stray molecules at the receiver. If *A_in_* exceeds a certain level of concentration (*λ*) in time slot *t_k_*, *Rx* interprets the received symbol as *sym*_1_ and *sym*_0_ otherwise. This process can be represented as

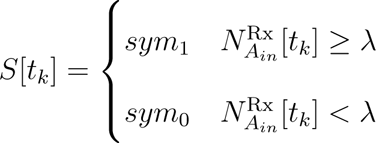

where *S*[*t_k_*] is the received or decoded symbol in the time slot *t_k_*.

## 4 Acknowledgements

## Graphical TOC Entry

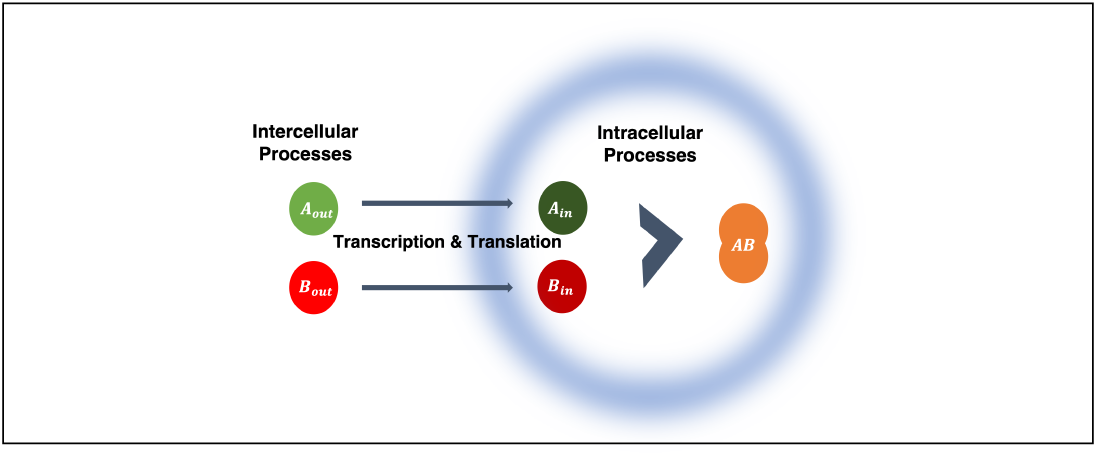

